# Parallelized calculation of permutation tests

**DOI:** 10.1101/2020.10.01.321828

**Authors:** Markus Ekvall, Michael Höhle, Lukas Käll

## Abstract

**Motivation:** Permutation tests offer a straight forward framework to assess the significance of differences in sample statistics. A significant advantage of permutation tests are the relatively few assumptions about the distribution of the test statistic are needed, as they rely on the assumption of exchangeability of the group labels. They have great value, as they allow a sensitivity analysis to determine the extent to which the assumed broad sample distribution of the test statistic applies. However, in this situation, permutation tests are rarely applied because the running time of naive implementations is too slow and grows exponentially with the sample size. Nevertheless, continued development in the 1980s introduced dynamic programming algorithms that compute exact permutation tests in polynomial time. Albeit this significant running time reduction, the exact test has not yet become one of the predominant statistical tests for medium sample size. Here, we propose a computational parallelization of one such dynamic programming-based permutation test, the Green algorithm, which makes the permutation test more attractive.

**Results:** Parallelization of the Green algorithm was found possible by nontrivial rearrangement of the structure of the algorithm. A speed-up – by orders of magnitude – is achievable by executing the parallelized algorithm on a GPU. We demonstrate that the execution time essentially becomes a non-issue for sample sizes, even as high as hundreds of samples. This improvement makes our method an attractive alternative to, e.g., the widely used asymptotic Mann-Whitney U-test.

**Availability:** In Python 3 code from the GitHub repository https://github.com/statisticalbiotechnology/parallelPermutationTest under an Apache 2.0 license.

**Supplementary information:** Supplementary data are available at *Bioinformatics* online.

## Introduction

Permutation tests are frequently used for non-parametric testing and are incredibly valuable within computational biology, with applications within genome-wide association studies [16, 1, 4], Pathway Analysis [20, 11], and expression quantitative trait loci studies [3, 21]. Monte Carlo-based sampling techniques [18] and exact tests that derive full permutation distributions are roughly the two ways to implement the permutation test.

Exact tests are traditionally seen as unattractive for large sample sizes, as the number of permutation grows super-exponentially with the sample size. Nonetheless, Green’s dynamic programming algorithm [7], which was made explicit by others [15, 22], partially overcome this computational problem. This algorithm is significant compared with any naive approach. However, the exact test’s popularity for larger sample sizes has not attracted much attention in the last couple of decades. We report here on an extension using a Graphics processing unit (GPU) implementation to compute parallelized exact tests and found it superior to the other tested alternatives in terms of speed and accuracy.

## Algorithm

Here we will describe our parallel Green method to calculate exact tests. First, in section, we describe the main objective, perform hypothesis testing with an exact test, then a description of the Green algorithm in section, and, finally, a description of how to parallelize the algorithm in section.

### Hypothesis testing

Let 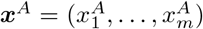 and 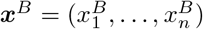 be two independent non-negative integers valued samples from the distributions 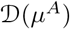 and 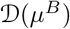 that only differ in their population means *μ^A^* and *μ^B^*, respectively. So, 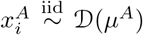, and 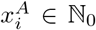 for 1 ≤ *i* ≤ *m*; and 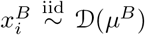, and 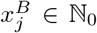 for 1 ≤ *j* ≤ *n*. We also form the concatenation of the samples as 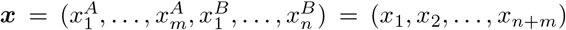. Our interest is in investigating the hypothesis *H*_0_ : *μ^A^* = *μ^B^* vs. the alternative *H_A_* : *μ^A^* > *μ^B^*. Note that we, without loss of generality, write the null-hypothesis as a point-hypothesis. If one instead has a composite null hypothesis of type *H*_0_ : *μ^A^* ≤ *μ^B^*, the procedure is to conduct the test under the most extreme parameter value of the null-hypothesis, which (for all relevant problems) is *μ^A^* = *μ^B^* [12, Sect. 5.9] and the test thus corresponds to the point-hypothesis case.

A way to perform the test is to determine how extreme the observed sum of sample **x**^A^, 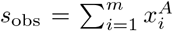 is under null hypothesis *H*_0_ : *μ^A^* = *μ^B^*. Typically, both null and alternative hypotheses assume a particular parametric family of distributions, parametrized by their respective mean parameters. From this, the test computes the *p* value, as the probability of observing *s_obs_* or a more extreme value in the alternative’s direction under the assumption that the null-hypothesis is true. However, it is rarely possible to analytically compute this probability, and one often has to resort to asymptotic approximations. A permutation test approach to the testing problem is to assume instead that the labels *A* and *B* are exchangeable under *H*_0_, as long as they stem from distributions only different in their mean parameters [10] and calculate how frequently samples with sample sums greater or equal than *s_obs_* appears when resampling from ***x***.

We can formulate the *p* value as Pr(*s_obs_* ≤ *S*|***x**, H*_0_), where Pr(*S*|***x**, H*_0_) is the probability mass function, and *S* is a random variable denoting the sum’s value in the first sample under the permutation distribution. The computationally expensive part is to obtain Pr(*S*) which is estimated by concatenating ***x**^A^* and ***x**^B^* to 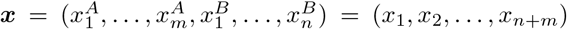, and draw all possible subsets of length *m* from ***x*** and count the number of occurrences of all possible sums; when the numbers of occurrences of all possible sums are available, the distribution is accessible.

Assume a random variable 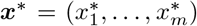, that is a randomly sampled subset from ***x*** with *j* elements and its corresponding sum is 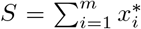, where *S* ∈ [0, *s*_max_]. Define *N*[*s*, *m*] to be the number of ways we can sample subsets ***x***\* with m elements in such a way that their sum *S* = *s*. Now Pr(*S* = *s*) can be expressed as the fraction of ways that a subset ***x***\* can be sampled so that its sum ends up to as *S* = *s* to the number of ways it can be sampled with any sum

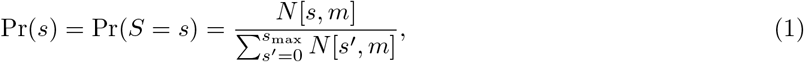

Combinatorics gives us that the denominator of the above equation 1 can be expressed as,

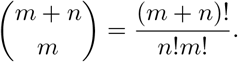

The calculation of the numerator is intricate. A naive algorithm that would exhaustively calculate the sum for each possible subset of size *m* and compare it to *s*, for all *s*, would need 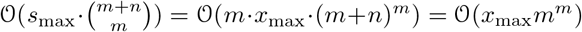 calculations, which becomes computational prohibitive even for moderate set sizes *m*. However, in section, we will discuss an algorithm solving the problem within polynomial time.

Now, the sought *p* value can be calculated as,

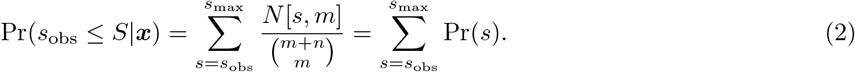

Alternatively, the same framework can be used to calculate the mid *p* value [17] as,

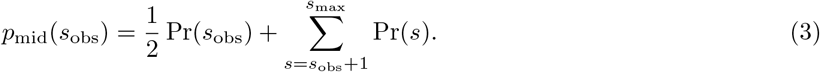

As mentioned above, there is no closed formula to calculate *N*[*s*, *m*]. However, it is possible to develop a dynamic programming algorithm to obtain *N*[*s*, *m*] [15, 22], described thoroughly in the next section.

### Efficient calculation of *N*[*s*, *m*]: the Green Algorithm

A dynamic programming algorithm for calculating *N*[*s*, *m*] was first presented in Green [7] and more in detail described by others [22, 6]. Here, we will give a walk-through of the algorithm, mostly to describe its parallelization in section. We also provide a simple use case of the algorithm in Supplementary Note 1.

We can find a recursive expression for *N*[*s*, *m*] by considering a scenario where ***x*\*** is drawn instead of the full set **x** from a subset 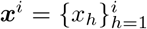 consisting of only the first *i* features of ***x***. To do so; we first need some definitions. Define *N_i_*[*s*, *j*] as the number of ways we can sample *j* elements so that their sum becomes *s* from a subset ***x^i^***. If we know how to calculate *N_i_*[*s*, *j*], we also know how to calculate *N*[*s*, *j*] = *N*_*i*=*m*+*n*_[*s*, *j*] = *N*[*s*, *j*], since 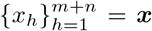. We also define *M_i_*[*s*, *j*] to be the number of ways we can sample ***x**^i^* so that the sample elements sum to *s* and including the last element *x_i_* in the sample.

We can now form a recursion of *N_i_*[*s*, *j*] as the number of ways to sample ***x**^i^* has to be equal to the number of ways to sample ***x***^*i*–1^ with the number of ways to sample ***x**^i^* that include *x_i_*. So,

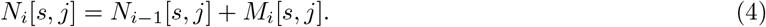

We can express *M_i_* in terms of *N*_*i*–1_ by noting that *M_i_*[*s*, *j*] = *N*_*i*–1_[*s* – *x_i_*, *j* – 1] when *x_i_* ≤ *s*, and otherwise zero. We can hence express the recursion 4 as,

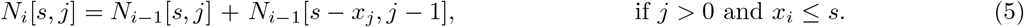

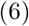

Let’s now turn to the boundary conditions of this recursion. The empty set 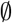 trivially reach the sum *s* = 0, thus

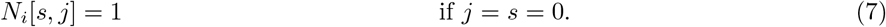

We can not sample *j* from *i* elements if *j* > *i*. Also, *N_i_*[*s*, *j*] has to be zero when either *i* ≤ 0 (i.e., checking the empty set of ***x***), when s < 0, or when *s*_max_ < *s* (i.e., when s is outside the boundary of possible sums). Hence,

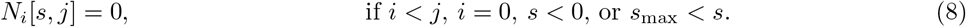

By combining the base case 8 and 7 with the generic sub-recursion 5, the final recursion is,

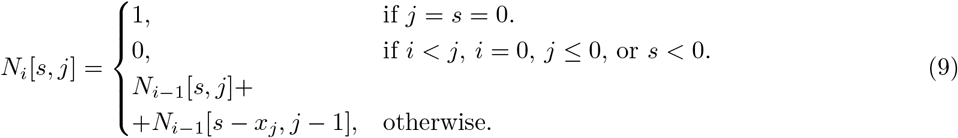

The pseudo-code of the top-down dynamic programming code of the recursion in Equation 9 is given in Algorithm S1. The algorithm needs some explanation. Instead of using one extensive array *N*[0… *s*_max_, 0 … (*m* + *n*), 0 … *m*], two smaller arrays, *N_old_*[0…*s*_max_, 0…*m*] and *N_new_* [0…*s*_max_, 0…*m*], are swapped and rewritten in each iteration of *i* in an oscillatory fashion — to save memory, see line 24. It is the complete rewriting of *N_new_* in the next iteration that makes this possible (any of the conditions in the recursion relation will re-calculate each entry *N_new_*).

We save quite some memory by keeping *N_new_* and *N_old_* instead of the full three-dimensional array. The former two arrays require 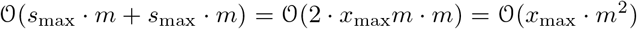, whereas the latter array requires 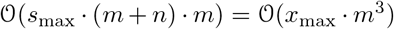. Moreover, this improvement in memory usage is a quintessential difference for the parallelized algorithm (GPU’s memory storage can easily be a bottleneck).

A second point, notice the structure of the for-loops; they could easily have been arranged in whatever way and still obtain correct computations. However, the dimensions of the two arrays have to be adjusted appropriately related to the outer-most loop. Nevertheless, this specific order has a purpose; it is parallelizable — described in the next section.

One final note on Algorithm S1, by plain observation on the nested loops, it easy to see that the running time is 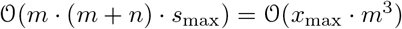.

### Parallelization over two dimensions: *s* and *j*

The meaning of parallelizing over two dimensions is, in this case, to fix one of the variables and check whether the two other variables are parallelizable given the third fixed variable. In practice, the fixed variable is the outer-most for-loop, and for each iteration of this variable, then within this loop, everything is calculated in parallel over the two other variables.

Out of the three variables, it is only necessary to find one such variable to fix, and instead of exhaustively checking all possibilities, one can check recursion 9 to see which variable is parallelizable. By comparing the left side to the right side, the only variable that is not dependent on contemporary action of itself is variable *i*, i.e.,
*N_i_*[*s*, *j*] = *f*(*s*, *s* – *x_j_*, *i* – 1, *j*, *j* – 1), and, furthermore, the other two variables are not possible to fix. Below is a verification that *N_i_*[*s*, *j*] is parallelizable given that *i* is fixed.

#### Initiation

In the lines 6 – 9 in algorithm S1, constants are set, and initialization of both arrays, *N_new_* [0…*s*_max_, 0…*m*] and *N_old_*[0…*s*_max_, 0…*m*], occur. There is no conflict for the parallelization of this step.

#### Maintenance

1 ≤ *i* ≤ (*m* + *n*) + 1

Consider iteration *i*. In lines 13 – 16, the array *N_new_* is only dependent on constants (i.e., the boundary conditions 8 and 7). Thus, the computation of *N_new_* is parallelizable. Furthermore, in lines 17 – 20, *N_new_* is only dependent on elements from *N_old_* and *x*, which both are invariant for *i* = 1 (except for *N_old_*, that switch values at the end of the iteration i, however, all computations for *i* are already done). Therefore, *N_new_* is parallelizable here too. Finally, at line 24, *N_old_* is switched with *N_new_*, and no parallelization occurs here. Hence, the algorithm is parallelizable for iteration *i*.

When entering the next iteration of *i*, i.e., *i* ← *i* +1, the same arguments above apply.

#### Termination

*i* = (*m* + *n*) + 2

When arriving at line 10 in algorithm S1 and *i* = (*m* + *n*) + 2, it will not enter the loop. By the maintenance of algorithm, one can be sure that the computation of *N*(0…*s*_max_, (*m* + *n*), *m*) is correct. Since there is no computation after the for-loop-block, hence, there are no more modifications on *N*(0…*s*_max_, (*m* + *n*), *m*), and it is safe to return this array.

### Discretization of real numbered samples into integer valued samples

As our test is defined for integer valued distributions of the samples, **x**. However, approximately we may use the procedure if we first discretize any real value distributed samples, ***y**^A^* and ***y**^B^*, where 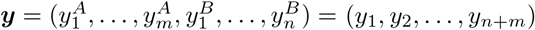. This was archived by partitioning the samples range into *n_w_* discretization windows, each of length, 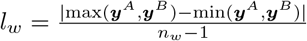. Each of these windows covers 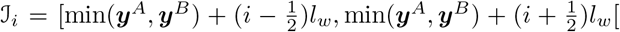, for *i* = 0; …, *n_w_* – 1, which enables us to map any continues sample into discrete variables, as 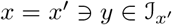, with values in the interval *x* ∈ [0, *n_w_* – 1]. Unfortunately, this comes to the cost of discretization errors, which will be a function of the number of discretization windows, *n_w_*. We will investigate the effects of such discretization in the Results sections below.

## System and methods

### Compared Methods

A couple of methods were used as comparison to our implementation. We implemented *t* tests through the function `scipy.stats.ttest_ind`, and Mann-Whitney *U* tests through `scipy.stats.mannwhitneyu`, both functions in scipy version 1.4.1. We downloaded the `FastPerm` method from the `master` branch of https://github.com/bdsegal/fastPerm. When executing `FastPerm` we applied the default parameter-tuning as described in the implementation guide, available at the method’s GitHub repository. We implemented the r-package Coin (https://github.com/cran/coin) version of the shift-method (exact test) as part of our python package and used it as a benchmark.

### System

Performance figures were recorded on a 4 core Intel i7-6700K with an NVIDIA GeForce RTX 2070 graphics card. Some of the experiments we compared this configuration’s performance to similar computers equipped with NVIDIA GeForce RTX 2060 and one with a NVIDIA Titan X Pascal graphics card.

### Implementation

A python 3.6 implementation implementing the below algorithms was made available under an Apache 2.0 license. We implemented our algorithm together with our discretization strategy as a CUDA [2] enabled Python module. The implementation and all results of this paper are available in reproducible form from a GitHub repository https://github.com/statisticalbiotechnology/parallelPermutationTest.

## Results

We implemented the algorithm described above, and set out to characterize the algorithm’s performance. To establish that the strategy executes in a practically useful time scale, we first tested the running time performance as a function of the number of discretization windows and its dependence on sample size. Subsequently, we tested the accuracy of our discretization strategy to establish that it is asymptotically unbiased and precise compared to other methods. Finally, we applied the method to a relatively large-scale proteomics dataset to establish the methods’ usefulness in a practical test scenario.

For some of the tests, we were able to benchmark the parallel Green algorithm against other methods. Here we implemented a single-thread version of the Green algorithm, downloaded a previously described Monte Carlo-based sampling method called the fast permutation method, `FastPerm` [18], the popular r-package `Coin` [9, 8] for exact permutation test, and used Python scipy’s implementation of the t test and Mann-Whitney *U* test.

### Test of running time requirements

#### Running time as a function of sample size, *n*

We first tested the running time requirements of the testes methods as a function of sample size. Here, we selected a uniform distributed samples, 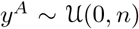 and 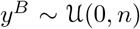, and drew samples of size *n* = |**y^A^**| = |**y^B^**| ∈ [50, 100, 200, 250, 300, 350, 400, 450, 500] = **n**. For each *n* ∈ **n**, using five samples replicates. The average time to calculate the corresponding *p* values was recorded and plotted as a function of **n** (Figure 1). For the tested sample sizes, the parallelized shift method scales well with sample size, and for all sample sizes, the parallelized shift method was found between 15 and 50 times faster than `FastPerm`, see Figure 1b at *n* = 300 and *n* = 100, respectively. However, they seem to reach a similar running time around *n* = 500.

**Figure 1:**
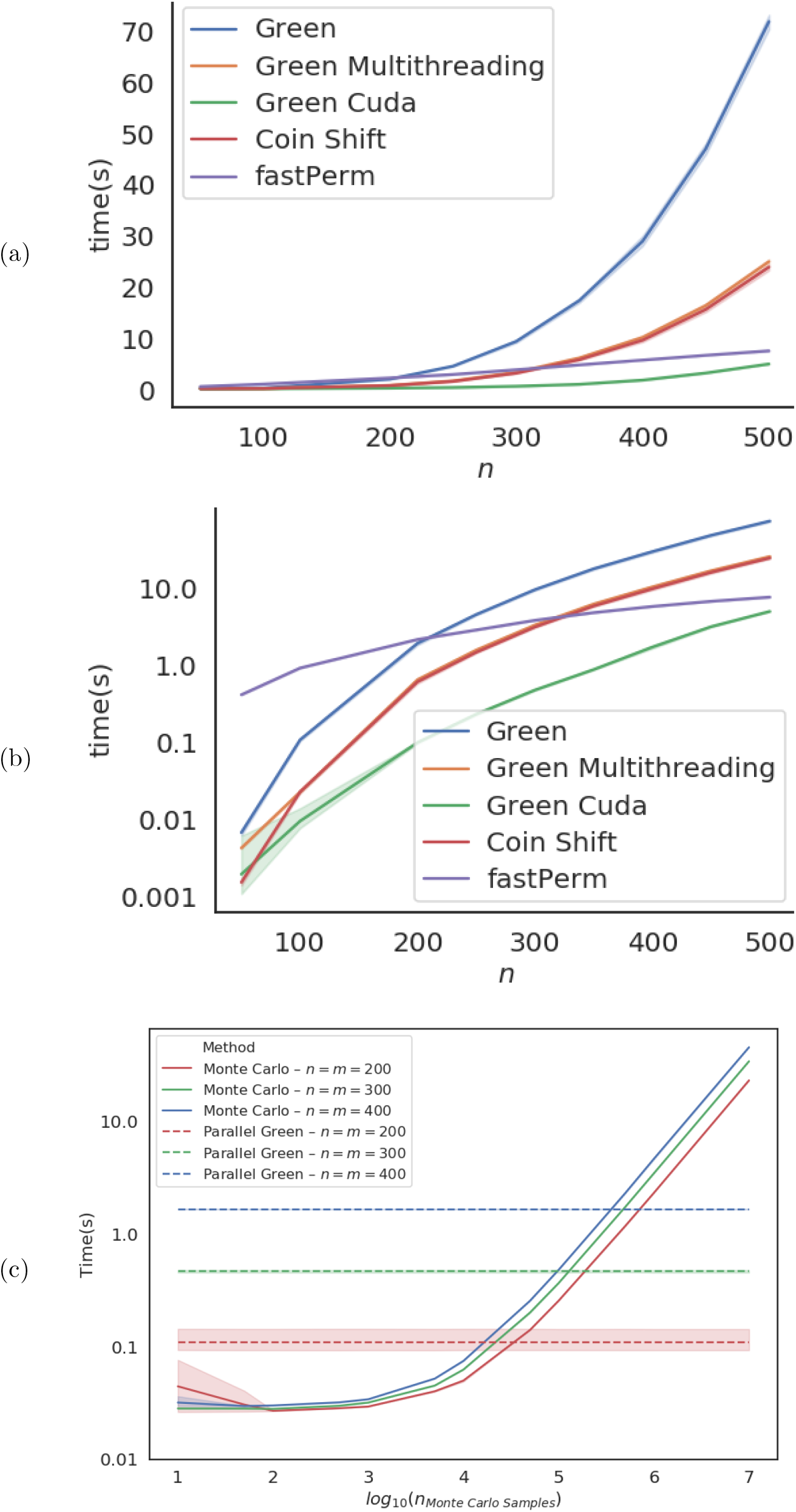
Running time requirements of the compared algorithms. We plotted the time required to calculate *p* values for samples from 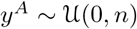 and 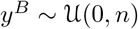 for different sample sizes **n**. The mean time and the 95% confidence interval around the mean time to calculate each of 5 replicate samplings was plotted in (a) linear and (b) log scale. It should be noted that the execution times where highly reproducible, and it might be hard to see the confidence interval in the plots. (c) We further invesigated the running time as a function of the number of Monte Carlo samples for a MC-based approach. We compared running time for a Monte Carlo sampled to draw samples of 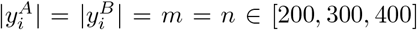, with five replicates. The horizontal lines displays the running time for parallel Green to compute the same samples.

The single-threaded Green algorithm should have a running time that expands as 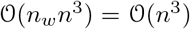 as *n_w_* was held constant in the experiment. This corresponds well with our observations in Figure 1.

Some Monte Carlo-based approaches are, unlike e.g. `FastPerm`, leaving the choice of the number of sampled permutations to their users. The number of permutations is inversely proportional the granularity of the *p* values estimated by the MC-sampler, and hence governs their possible precision, at least when no other techniques such as importance sampling is used. We hence measure the running time for different amounts of permutation samples with the MC-sampling functionality of the package `Coin`, putting it in context of the running time of Parallel Green (Figure 1c). The two methods seem to have similar running times for about 10^5^–10^6^ permutation samples, tentatively suggesting that the parallel Green will be the faster method of the two methods, whenever a precision of the estimated *p* values is desired to be better than 10^−5^–10^−6^.

Furthermore, to see how the running time depends on the hardware, the test was repeated on other GPUs. However, we found relatively small difference in performance between the tested graphic cards (see Supplementary Figure S1.

#### Running time as a function of the number discretization windows, *n_w_*

We also wanted to establish that our implementation scales well with the number of discretization windows used for the test. Again, we sampled from 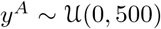 and 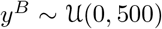 with sample size *n* = *m* = 500, with five replicates. We plotted the running time as a function of *n_w_* ∈ **n_w_** = [50,100, 200, 250, 300, 350, 400, 450, 500] in Figure 2.

**Figure 2:**
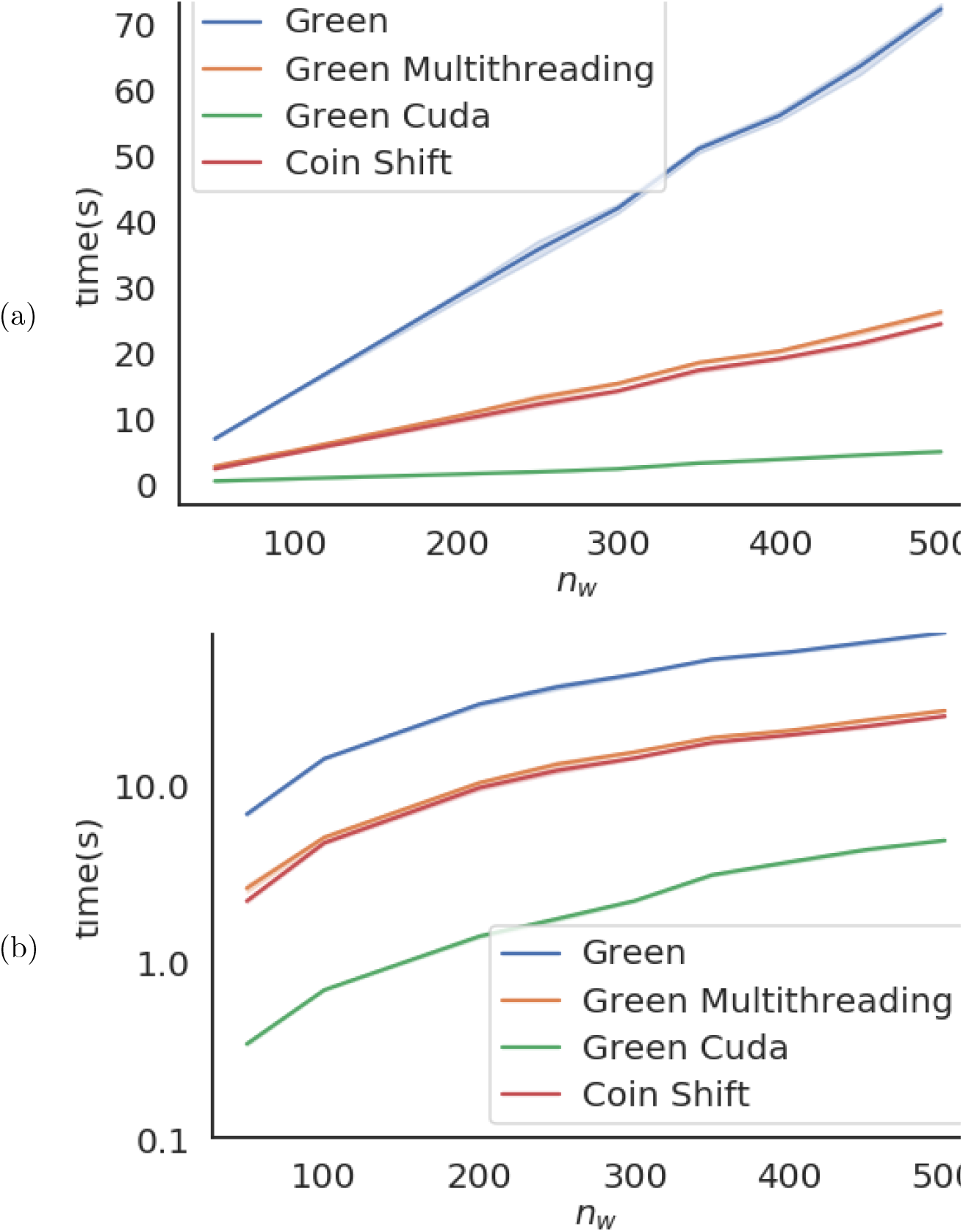
Running time as a function of the number of discretization windows of single-thread and parallel Green algorithm. We plotted the mean of the mean and standard deviation of the required running time, as wall time, as a function of the number of discretization windows, *n_w_*. Time was plotted (a) in normal scale, and (b) log-scale. Note that the other methods were excluded from the plot, as the discretization step is exclusively present in the Green algorithm.

The single-threaded implementation of the Green algorithm should theoretically have a running time complexity 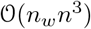. As sample size *n* is kept constant in the experiment, we expect the running time to expend as 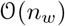. Figure 2 confirms this for the single thread Green.

### Memory Allocation

#### Test of Memory allocation

We characterized the memory requirements of parallel Green by increasing sample size *n* and *m*, and a growing number of bins *n_w_*. The first experiment (Figure 3(a)), varied the set sizes |**y**^*A*^| = |**y**^*B*^| = *m* = *n* ∈ **n** = [10, 20, 30,…, 480, 490, 500], for three different bin sizes *n_w_* of 64, 128, and 256. In the second experiment (Figure 3(b)), the number of bins was the variable *n_w_* ∈ **n_w_** = [10, 12, 14,…, 396, 398, 400] for three different set sizes |**y**^*A*^| = |**y**^*B*^| = *n* = *m* of 125, 250, and 500. In both experiments, the data was sampled from 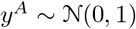 and 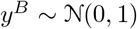, and the number replicates was 1000.

**Figure 3:**
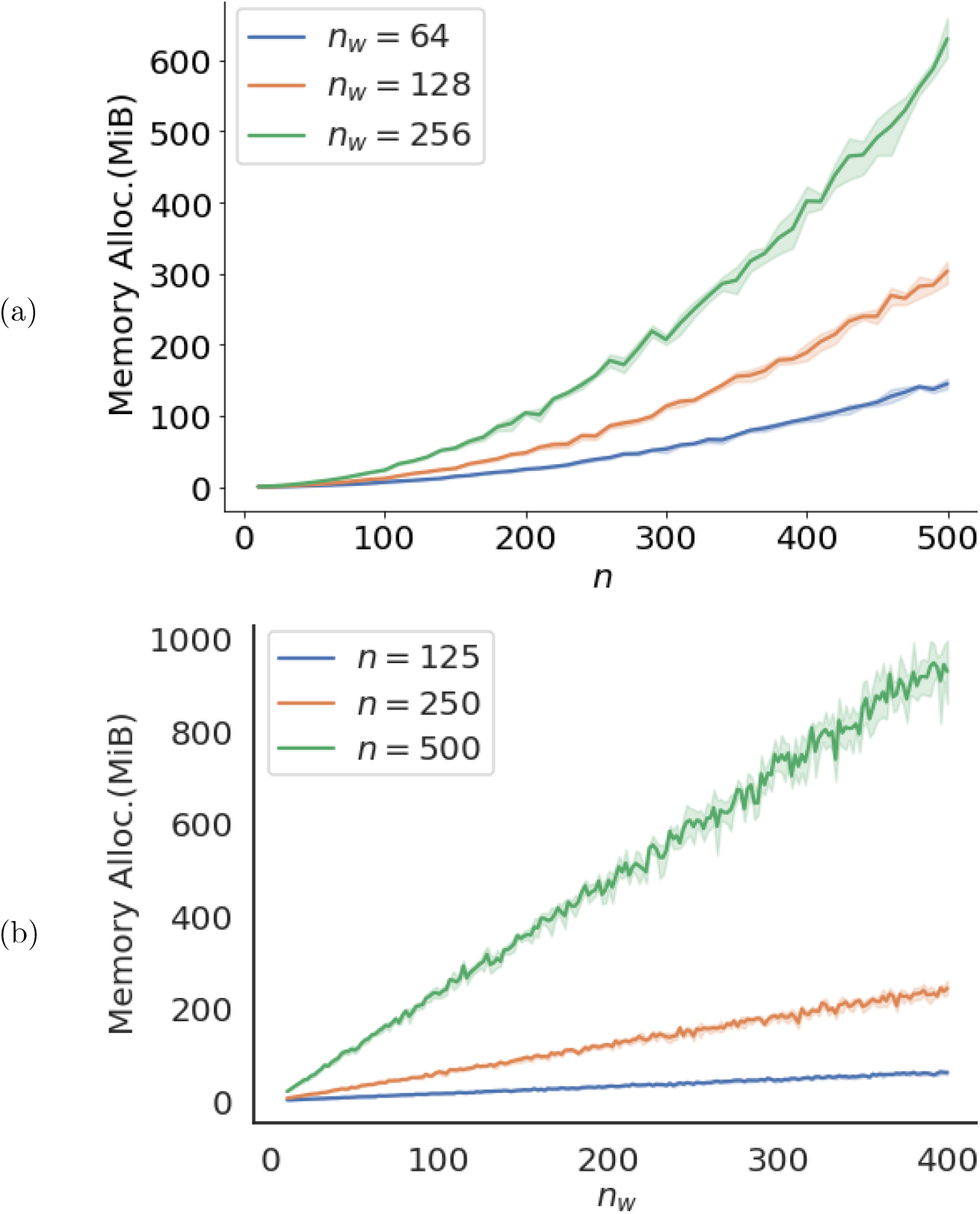
Memory allocation as a function of set size and bin size. We plotted the required memory allocation to calculate *p* values for samples from 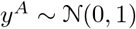 and 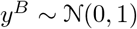 for different sample sizes **n** and bin sizes **n_w_**. In figure (a) the different *n_w_* used were 64,128, and 512 with 5 replicates, and in figure (b) the different *n* were 125, 250, and 500 with 5 replicates.

The amount of data that can be handled by a GPU at each point in time is limited by the GPU’s memory size. Here we used an NVIDIA GeForce RTX 2070, which allows for 7982 MiB memory allocation. One could imagine settings where s_max_ is so large that one can not calculate *N* even for one data point. However, we would instead run into floating-point problems before reaching this type of memory problem for such cases. For large set-sizes, 1000 < |**y^A^**| + |**y^B^**|, the sums in the entries in *N* start go beyond the maximum double-point precision in CUDA i.e., ≈ 1.79*e*^308^. A future improvement of our algorithm could be to reduce the values in each iteration by dividing all elements in *N_i_*[*s, j*] by a normalizing factor in each iteration over *i*, which would help us reduce the problems of *N* becoming too large.

### Test of accuracy and precision

#### Test dependency of window size

While the Green Algorithm is a non-parametric test for discrete test statistics, the algorithm’s performance will be a function of the number of windows, *n_w_*, we use for discretizing any continuous data. We hence wanted to characterize the influence of *n_w_* on the accuracy of our test. We selected samples from a Normal distribution and compared the computed *p* values with the ones from a regular *t* test (Supplementary Figure S2). The results suggest that both the accuracy and precision of the test improves when increasing the number of windows. However, the effect seems to saturate for *n_w_* > 30.

#### Comparison to other method’s accuracy and precision against comparative methods

We subsequently wanted to test the accuracy of the estimated *p* values. Again we used Normal distributed samples and used *t* test-calculated *p* values as reference. As benchmark comparisons, we again used the `FastPerm` method and Python’s asymptotic Normal distribution implementation of the Mann-Whitney *U* test, `scipy.stats.mannwhitneyu`. We plotted the ratio, 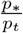, where *p*_*_ is the tested *p* value and *p_t_* is given by a *t* test, as a function of sample size for two different effect sizes (Figure 4). Overall, the 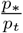 of the parallel Green were found closer to 1, and less dependent on the sample size than the ones from the compared methods. The reason for the Mann-Whitney *U* test deviating from the results of the *t* test, particularly in Figure 4b, is that the test has comparatively low efficiency when testing on normal distributed data. It is also important to note that the Mann-Whitney *U* test, which depends on ranking statistics, would not be under the same null model as the compared methods in the presence of ties. However, we expect ties to be rare when sampling from continuous distributions.

**Figure 4:**
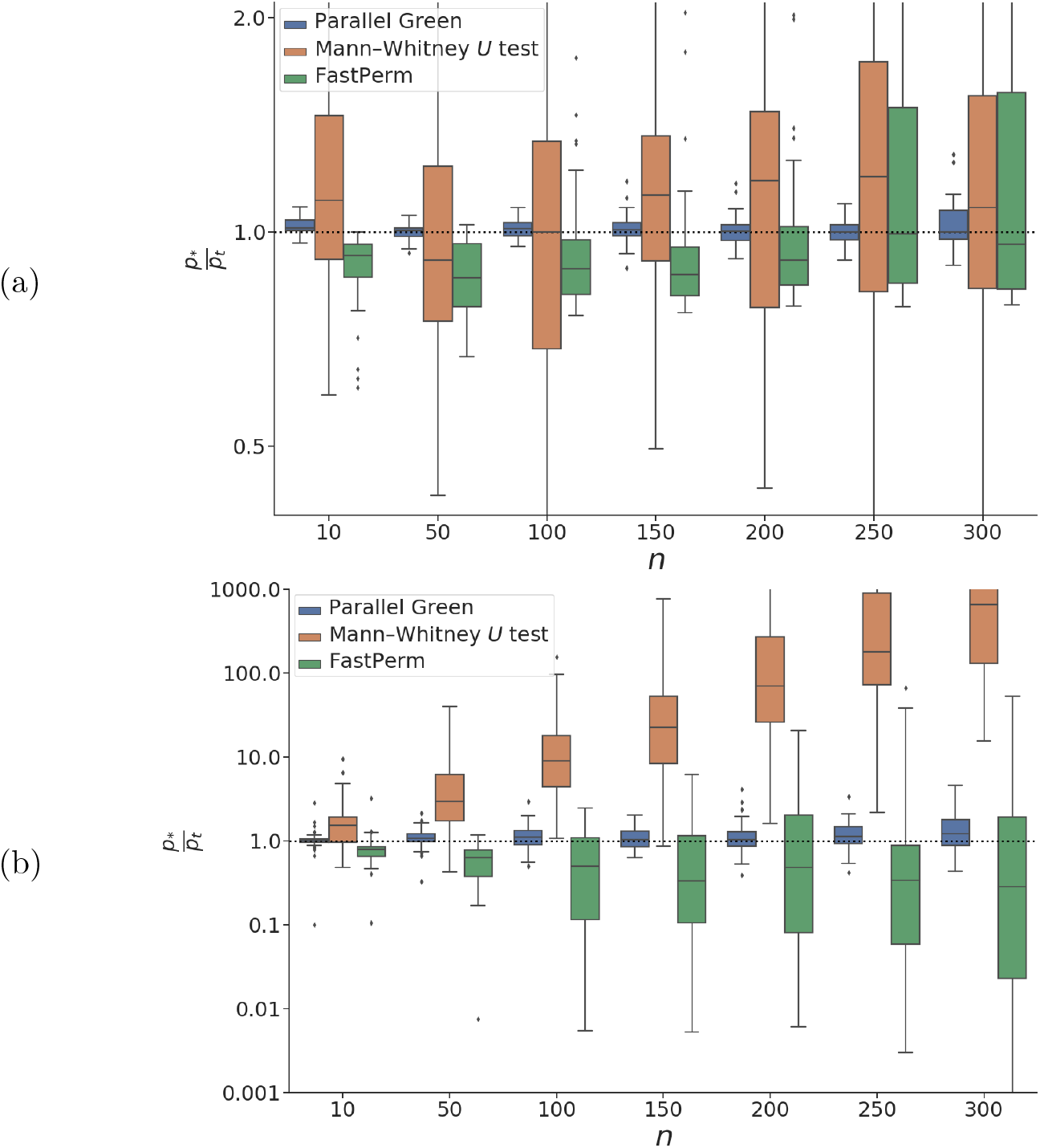
Comparison of estimation error as a function of sample size. We plotted the fold change between one side parallel Green, FastPerm and Mann-Whithey *U* test and on the other side a *t* test, as a function of the sample size, *n*, when 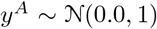 and (a) 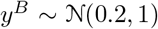, and (b) 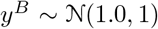. For both cases we plotted results from 50 samples, and used *n_w_* = 100 discretization windows.

#### Calibration test

We also tested the parallelized shift method’s *p* values uniformity under the null hypothesis [14]. This property is of particular importance for studies where we test many variables for the same sample, as it is the base for efficient multiple hypothesis testing [19, 5]. Here, we compared parallel Green against the FastPerm method, Mann-Whitney *U* test, and a regular *t* test. We picked 10000 samples from the Normal distribution and a log-Normal distribution and plotted each method has estimated *p* values as a function of their relative rank (Supplementary Figure S3). We found that the calibration of the parallel Green method was on par with the *t*est. However, unsurprisingly the calibration seems to be entirely off for the t tests on log-Normal distributed data. We see that the Mann-Whitney *U* test’s calibration appears conservative, while the FastPerm method appears anti-conservative for both tested distributions.

### Running time requirements for a Proteomics dataset

As the last test, we tested the algorithm’s performance on a dataset of breast cancer samples from the CPTAC consortium [13]. For the 8051 proteins for which measurements had been obtained for all samples, we tested differential abundance between 80 non-triple-negative and 26 triple-negative samples. The run-time for parallel Green method can be found in table 1. For the other methods: `FastPerm` took 45 minutes, 1.5 seconds for a Mann-Whitney *U* test, and 1.32 seconds for a *t* test.

**Table 1:** Running time for parallel Green on the proteomics dataset as a function of the number of discretization windows *n_w_*

| $n_w$ | 16 | 64 | 256 | 512 | 1028 |
| --- | --- | --- | --- | --- | --- |
| $time(s)$ | 6,31 | 6,87 | 10,8 | 17,5 | 37 |

## Discussion

Statistical testing is the base for most scientific activities. Also, in most research areas, the amount of public data is rapidly increasing, and hence there is a need for ever more efficient methods to compute significance. Permutation tests offer an exciting method as they do not assume a particular sampling distribution, but instead, build one by permuting label associations to the observed data. This approach corresponds perfectly with the null hypothesis that there is no difference in case and control outcomes.

Here, we have described a parallelized dynamic programming method to perform permutation tests. We have demonstrated that it is faster and more accurate than the sampling-based methods. Previous work by Pagano and Trichler [15] demonstrates that one can quickly expand exact tests to handle missing values, something that rank-based not easily can handle. We note that several studies are dependent on normal approximations of non-parametric tests such as the Mann-Whitney *U* test. In practice, the implementation of such tests is approximations as they are asymptotic and not exact. The parallel Green method offers an exact test that does not appear much slower but more accurate than such tests. However, admittedly, the difference between the outcomes of asymptotic and permutation tests gets smaller with increased sample size.

Permutation tests have been successfully used in many, if not most areas of bioinformatics as a relatively assumption free method for assessing statistical significance in inferences. In most applications the procedures involve some flavor of Monte Carlo-based sampling methodology. Here we demonstrated that for at least in the case of statistical hypothesis testing on can instead rely on calculating a full distribution of the sampling space.

## Acknowledgements

This research was supported by grants from the Swedish Research Council, (2017-04030) and Swedish Foundation for Strategic Research (BD15-0043), as well as a donation of the Titan X Pascal GPU used for this research from the NVIDIA Corporation.

