## Supplementary Material for "Parallelized calculation of permutation tests"

**Algorithm S1** # cardinality-$j$ subsets s.t. their elements sum is equal to $s$.

```
m: Scalar equal to the number of samples in x^A.
n: Scalar equal to the number of samples x^B.
s_max: Scalar equal to the sum of the m largest values in x.
x: Concatenated vector of x^A and x^B with length m + n.
procedure NumberOfSubsets(m, n, s_max, x)
    x = [0] + x                                      ▷ Include the empty set ∅ to x
    m = m + 1                                        ▷ Correct the indices for this inclusion.
    Allocate array N_old[0...s_max, 0...m] with zeros
    Allocate array N_new[0...s_max, 0...m] with zeros
    for i = 1 to (m + n) + 1 do
        for j = 1 to m + 1 do
            for s = 0 to s_max + 1 do
                if s = (j − 1) = 0 then                  ▷ Sub-recursion 7
                    N_new[s, j − 1] = 1
                else if i < j then                       ▷ Sub-recursion 8
                    N_new[s, j − 1] = 0
                else if j > 1 and x[i − 1] ≤ s then      ▷ Sub-recursion 5
                    N_new[s, j − 1] = N_old[s − x[i − 1], j − 2] + N_old[s, j − 1]
                else if j > 1 and x[i − 1] > s then
                    N_new[s, j − 1] = N_old[s, j − 1]
                end if
            end for
        end for
        N_old = N_new;                               ▷ Update N_old for next iteration.
    end for
    return N_new[0...s_max, m]                       ▷ By remark ??: N_{m+n}[s, m] = N(s, m)
end procedure
```

(a)

(b)

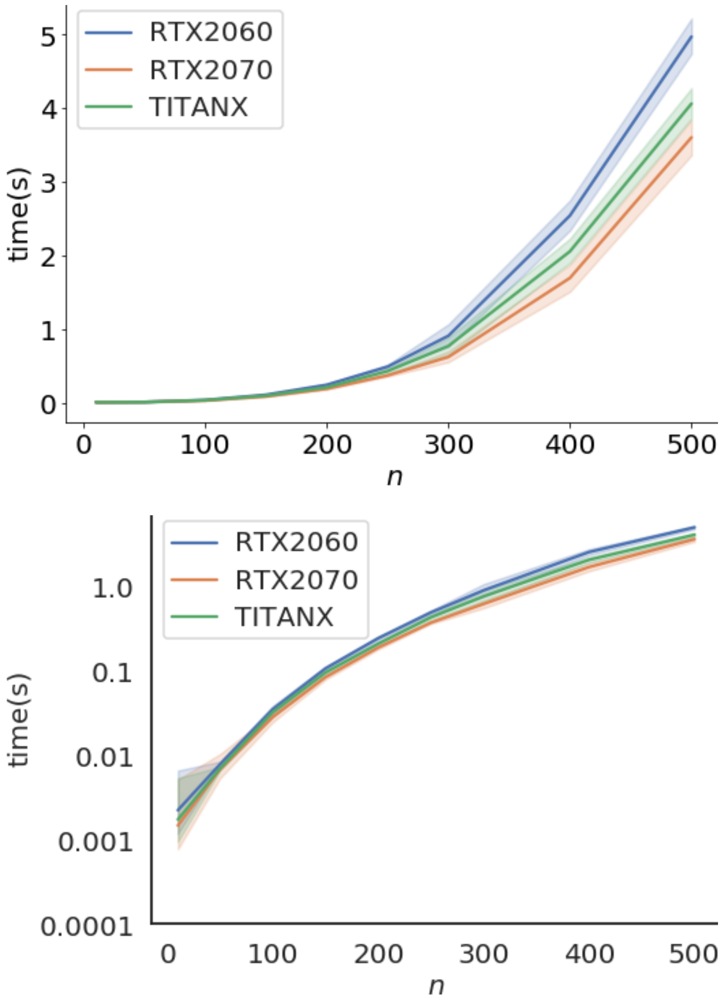


Figure S1: **Run time requirements of different GPUs**. We plotted the time required to calculate $p$ values for samples from $y^A \sim \mathcal{N}(0.2, 1)$ and $y^B \sim \mathcal{N}(0.2, 1)$ for different sample sizes $\mathbf{n}$ on similar computers with different graphics cards. The mean time and the 95% confidence interval around the mean time to calculate each of 5 replicate samplings was plotted in (a) linear and (b) log scale. It should be noted that the execution times where highly reproducible, and it might be hard to see the confidence interval in the plots.

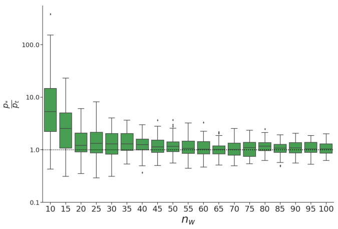


(a)

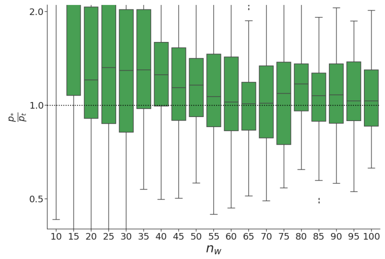


(b)

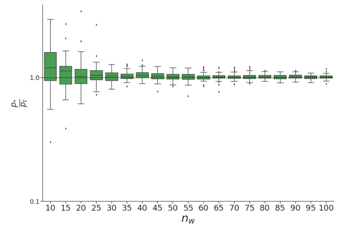


(c)

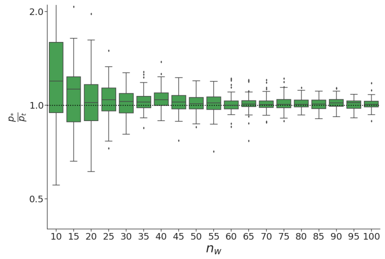


(d)

Figure S2: **The influence of $n_w$ on the accuracy and precision of the Green algorithm.** We plotted the fold change difference between the $p$ values calculated by the Green algorithm and a $t$ test, as a function of the window size, $n_w$. Here we sampled 50 samples with sizes $|y^A| = |y^B| = 150$ from each of $y^A \sim \mathcal{N}(0.0, 1)$ and (a)(b) $y^B \sim \mathcal{N}(1.0, 1)$, or (c)(d) $y^B \sim \mathcal{N}(0.2, 1)$. Here we selected window sizes were $n_w \in \mathbf{n_w} = [10, 15, \ldots, 95, 100]$. (a)(c) We plotted the full range and (b)(d) details of $\frac{p_*}{p_t}$ in the range $[0.4 - 2.1]$.

(a)

(b)

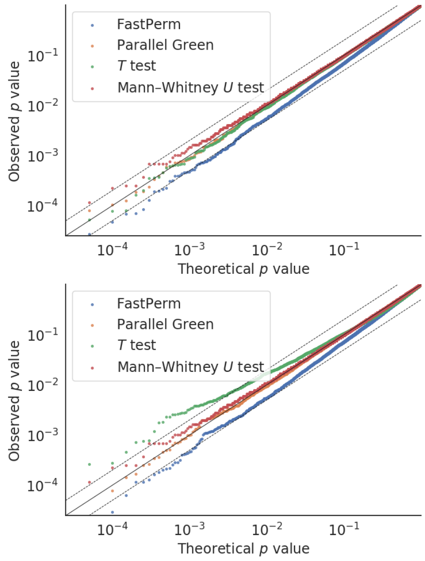


Figure S3: **Calibration plot, i.e. test of $p$ values uniformity under null.** We compared the parallelized shift method, the Mann-Whitney $U$-test, and a $t$-tests uniformity under null. We tested 20000 samples with (a) $y_i^A, y_i^B \overset{\text{iid}}{\sim} \mathcal{N}(0,1)$ and (b) $y_i^A, y_i^B \overset{\text{iid}}{\sim} \log\mathcal{N}(0,1)$. In both cases the sample sizes were set to $|y_i^A| = |y_i^B| = 20$ and the number of discretization windows, $n_w$, were set to, 100. The continuous line $y = x$, and the dashed lines $y = 2x$ and $y = x/2$ were included as reference.
